# Cambium-specific Transcriptome Analysis of Paulownia to Study the Molecular Impacts of Winter and Spring Seasons on Tree Growth

**DOI:** 10.1101/2020.10.27.357582

**Authors:** Zachary D Perry, Thangasamy Saminathan, Alok Arun, Brajesh N Vaidya, Chhandak Basu, Umesh K Reddy, Nirmal Joshee

## Abstract

Paulownia (*Paulownia elongata*) is a fast-growing, multipurpose deciduous hardwood species that grows in a wide range of temperatures from –30 °C to 45 °C. Seasonal cues influence the secondary growth of tree stems, including cambial activity, wood chemistry, and transition to latewood formation. In this study, a *de novo* transcriptome approach was conducted to identify the transcripts expressed in vascular cambial tissue from senescent winter and actively growing spring seasons. Illumina paired-end sequenced cambial transcriptome generated 297,049,842 clean reads which finally yielded 61,639 annotated unigenes. Based on non-redundant protein database analyses, Paulownia cambial unigenes shared highest homology (64.8%) with *Erythranthe guttata*. A total of 35,471 unigenes resulted from KEGG annotation that were mapped to 128 pathways with metabolic pathways dominated among all. Additionally, DEG analysis showed that 2,688 and 7,411 genes were significantly upregulated and downregulated, respectively in spring compared to winter. Interestingly, quite a number of transcripts belonging to heat shock proteins were upregulated in spring season. RT-qPCR expression results of fifteen wood-forming candidate genes involved in hemicellulose, cellulose, lignin, auxin and cytokinin pathways showed that the hemicellulose genes (*CSLC4, FUT1, AXY4, GATL1*, and *IRX19*) were significantly upregulated in spring season tissues when compared to winter tissues. In contrast, lignin pathway genes *CCR1* and *CAD1* were upregulated in winter cambium. Finally, a transcriptome-wide marker analysis identified 11,338 Simple Sequence Repeat (SSRs). The AG/CT dinucleotide repeat predominately represented all SSRs. Altogether, the cambial transcriptomic analysis reported here highlights the molecular events of wood formation during winter and spring. The identification of candidate genes involved in the cambial growth provides a roadmap of wood formation in Paulownia and other trees for the seasonal growth variation.

## Introduction

Paulownia (*Paulownia elongata*) is an extremely fast-growing woody plant reaching up to 20 feet in one year when young. Some species of *Paulownia* when in plantation can be harvested for saw timber in as little as five years. The genus *Paulownia* consists of nine species of deciduous, fast growing, multi-purpose, and hardwood trees (Zhu *et al*., 1986) and have long been shown to be extremely adaptive to wide environmental variations in both edaphic and climatic factors, as well as being capable of growing on marginal lands (Clatterbuck and Hodges, 2004; Sedeer and Nabil, 2003). *Paulownia* species are native to Asia from China, Laos, and Vietnam and grown in Japan and Korea. It has been cultivated in Australia, Europe, and both North and Central America. Ten-year-old Paulownia tree in natural conditions can attain 30–40 cm in diameter at breast height (DBH) and a timber volume of 0.3–0.5 m^3^ (Zhu *et al*., 1986). Craftsman in Japan and other countries have used this valuable wood to create intricate carvings, surfboards, musical instruments, toys, and furniture. Paulownia wood has a high ignition point of 420–430 °C compared to other hardwoods which range generally from 220–225 °C (Akyildiz and Kol Sahin, 2010). The wood of Paulownia has also been shown to be fire retardant (Li and Oda, 2007) as it burns at much higher temperature in comparison to many other wood species. Paulownia bears abundant flowers that are highly nectariferous and yield premium honey (Yadav *et al*., 2013) adding to the rural economy. By adding Paulownia wood flour (25–40%) to plastics, an attractive, equally strong, environmentally agreeable, and economically important biocomposite can be produced (Tisserat *et al*., 2013a; Tisserat *et al*., 2013b, 2015) that can serve many industries. In addition, due its light weight and strong nature of wood, it is attracted by music industry to make soundboards of stringed musical instruments such as the guqin, guzheng, pipa, koto, gayageum and electric guitars. Biochar produced from Paulownia is also a desirable organic soil amendment which allows the growth of beneficial microbes in the porous holes of the biochar (Vaughn *et al*., 2015). Recently, researchers found its potential use as an animal feed resource (Stewart *et al*., 2018).

Wood synthesis provides one of the most important sinks for atmospheric carbon dioxide (Ye and Zhong, 2015). Wood formation is a result of the regulated accumulation of secondary xylem cells (fibers, vessels, and rays in dicots) differentiated from the vascular cambium that involves wall thickening. This wall thickening is accompanied by the biosynthesis of wall components, lignin, cellulose, and hemicelluloses, and it is terminated by programmed cell death (Samuels *et al*., 2006; Song *et al*., 2006). In order to survive for multiple growing seasons, perennial plant species have adapted a dormancy regulation system which allows active growth during the desirable time of year, and vegetative dormancy when climatic conditions are unfavorable for growth (Shim *et al*., 2014). Being one of the fastest growing tree species, a Paulownia tree is capable of producing ∼45 kg/tree in the first growing year and ∼90 kg/tree at the end of second year (Joshee, 2012). Paulownia being a perennial tree, harvest is not limited to a small seasonal window but can be conducted year-round with proper management practices. Another beneficial property of Paulownia is coppicing, the production of multiple sprouts from a stump after the removal of the tree or shrub. Harvest cycles of 2–3 years could be implemented to establish a short rotation fast growing bioenergy crop. Since Paulownia is a short rotation and fast-growing perennial tree, it serves as a good candidate for the production of lignocellulosic biofuel which can eliminate dependence on fossil fuel. Further, trees may also have a benefit as stable wildlife habitat because they are not disturbed by annual harvests.

*Arabidopsis thaliana* has been used widely as a model system for secondary growth. However, the drawback to using *Arabidopsis* is the fact that it is an herbaceous plant, lacking secondary growth. In order to combat this feature, mutant lines were developed which have phenotypes exhibiting secondary growth characteristics. Initial studies used *A. thaliana* microarrays to determine the differential expression of transcripts related to secondary growth (Ko and Han, 2004; Oh *et al*., 2003). The next generation of studies utilized expressed sequence tags (ESTs) to determine a genomic “snap shot” of how wood is formed (Moreau *et al*., 2005; Schrader *et al*., 2004). Transcriptome analyses of various tree species indicate involvement of receptor kinases, transcription factors, and secondary wall biosynthesis genes that are highly expressed in wood-forming cells (Aspeborg *et al*., 2005; Dharmawardhana *et al*., 2010; Pavy *et al*., 2008; Wang *et al*., 2009; Wilkins *et al*., 2009).

In the recent past, scientific research addressing Paulownia transcriptomics have been accumulated. However, focus of the studies has been on drought tolerance, and the analysis of a phytoplasma that causes Witches Broom Disease (Dong *et al*., 2014a, b; Mou *et al*., 2013). Comparative analysis of microRNA expression (Cao *et al*., 2018a), regulation of long noncoding RNAs (Fan *et al*., 2018), and genome-wide analysis of lncRNAs (Cao *et al*., 2018b) provided comprehensive transcriptome analyses with *phytoplasma* infection. Transcriptome sequencing and a *de novo* assembly approach were together used to analyze gene expression profiles in *P. fortunei* infected by *Phytoplasmas* (Fan *et al*., 2014). Studies have been carried out to analyze the variations between *Paulownia tomentosa* and its autotetraploid counterpart to characterize the differential expression of unigenes (Fan *et al*., 2015) and microRNA expression under drought stress (Zhao *et al*., 2018). An investigation into the physiological alterations of *P. fortunei* x *P. tomentosa* in response to infection by Paulownia witches’ broom (PaWB) (*Phytoplasma spp*.), a pathogenic bacteria responsible for crop losses worldwide, was conducted by differential expression analysis of RNA-seq data of infected vs. pathogen free specimens (Mou *et al*., 2013). Another study investigated the expression of unigenes derived from Witches’ broom infected *P. tomentosa* x *P. fortunei* by a *De novo* assembled transcriptome (Liu *et al*., 2013). Recently, transcriptome and small RNA sequencing analysis revealed roles of PaWB-related miRNAs and genes in *Paulownia fortunei* (Li *et al*., 2018). An investigation into the miRNAs related to the regulation of gene expression in both *P. australis* diploid and autotetraploid genotypes was performed by sequencing of small RNA libraries for the two respective genotypes (Niu *et al*., 2014). Experimentation by (Li *et al*., 2014) identified the genes related to a synthesized autotetraploid of *P. tomentosa* x *P. fortunei*, which exhibits advanced characteristics such as greater yield and higher resistance than the diploid wild type tree.

Cambial development, the initiation and activity of the vascular cambium, leads to an accumulation of wood, the secondary xylem tissue. Seasonal cues play a significant role in determining cambial growth as perennial plants growing in temperate and high-latitude regions show termination of cell division in the meristems (Nitsch, 1957) and reversal of growth arrest during long days (Espinosa□Ruiz *et al*., 2004). Time-coursed transcriptome analysis identified participation and modulation of hormone-related genes; IAA, ARF and SAURs were downregulated and circadian genes including PIF3 and PRR5 were upregulated (Wang *et al*., 2019). Transcriptome data from the same tissue/s at different time points or of different physiological conditions were compared to one another to elucidate the gene expression pattern to each treatment. Interconnected signaling profiles between cytokinin and auxin indicated that they displayed distinct distribution across the cambium with increased cytokinin content to stimulate cell division (Immanen *et al*., 2016). Interestingly, cambial zone in addition to showing elevated cambial cytokinin level, the cambial auxin concentration and auxin-responsive gene expression were also increased. In Paulownia, by investigating the differential expression of vascular cambium, the site of lateral growth and xylem production (Nieminen *et al*., 2015), tissue during wood-forming spring and senescent fall seasons, the transcripts which influence the production of wood in *P. elongata* were determined. Recent study provided evidence for the involvement of microRNAs in *Paulownia tomentosa* cambial tissues in response to seasonal winter and summer changes (Qiu *et al*., 2018).

Transcriptomic analyses have been carried out to profile gene expression regulations for biotic and abiotic stresses, and growth responses. However, to the best of our knowledge no study has described how the gene expression profile changes in woody tissues under seasonal variations. In this study, we sequenced and analyzed the transcriptomes of cambium tissues collected during winter and spring seasons to assess the impact of two seasons on biomass. A transcriptome-wide analysis identified 61,639 annotated unigenes, and 2,688 and 7,411 transcripts were up- and downregulated, respectively in spring season. Interestingly, among selected wood-forming genes, hemicellulose-specific genes were upregulated in spring. Finally, 11,338 Simple Sequence Repeat (SSRs) were identified from the transcriptome data. The identification of genes and pathways involved in cambial growth will be useful to further investigate the regulation of wood formation in Paulownia and other trees.

## Materials and Methods

### Collection of Cambium Tissues and RNA Isolation

Samples were randomly selected from trees at the FVSU Paulownia bioenergy plot located at 32° 31’15.04” N and 83° 52’12.95” W. Samples were taken in triplicates during two two seasonal points, each seasonal point representing a different physiologic state. The first sample (Winter Wood – hereafter referred as WW) was collected in March, 2015 and represented the senescent winter wood (Figure 1A). The second sample (Spring Wood – hereafter referred as SW) was collected in May, 2015 and was representative of the actively growing spring wood (Figure 1B). Samples were harvested from twigs which were located at a height of 1.0–1.5 m from the ground, and having a diameter of 2.0–3.0 cm. Since Paulownia has an opposite branching pattern, the Spring and Winter samples were taken from the same node positioned on opposite sides. Sections of limb (15–30 cm long) were collected by removing the selected limb section with an ethanol (70 %) solution and RNAse*Zap®* (Ambion, Foster City, CA, USA) treated pruner and gloved hands. The samples were labeled, wrapped in aluminum foil, flash frozen in liquid nitrogen, and subsequently stored in a -80 °C freezer until further use. Biological replicates were labelled as WW1, WW2 and WW3 for winter, and SW1, SW2, and SW3 for spring, respectively.

**Figure 1.**
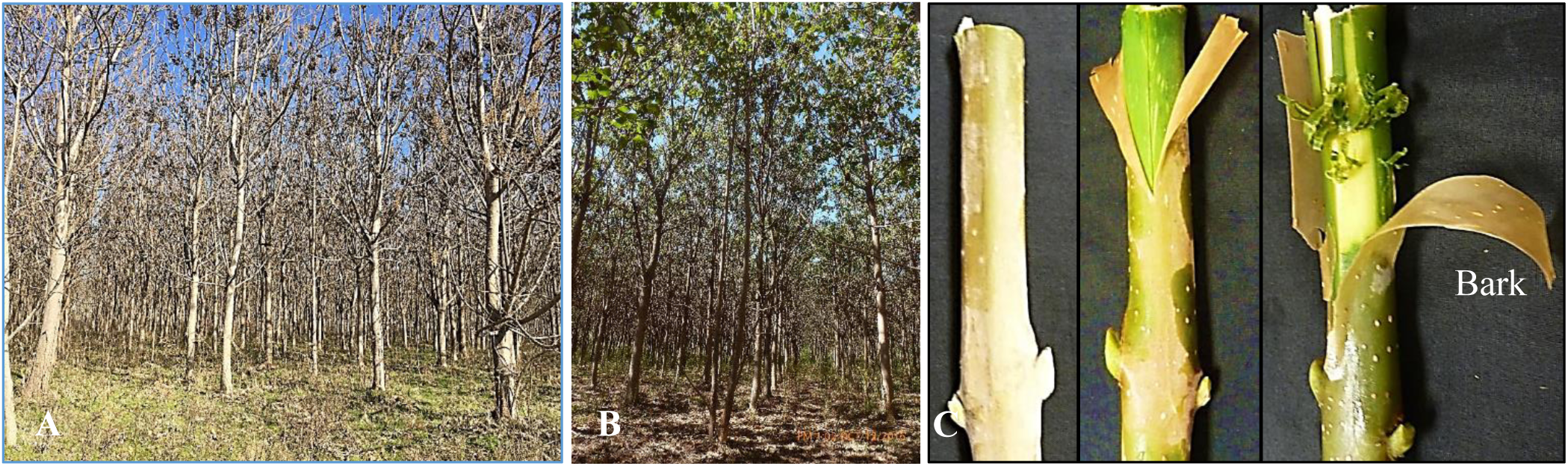
Paulownia tree growth and sampling of tree twigs. Paulownia Demonstration Plot at FVSU showing dormant during winter (A) and active growth during spring (B). Cambial tissue sampling from paulownia twig (C); unopened twig (left), bark removal (middle) and scrapping wood forming cambial tissue for RNA extraction (right).

For high quality intact total RNA extraction, vascular cambium tissues were harvested from the frozen samples by first slicing a shallow, longitudinal, cut into the outer bark with a sterile scalpel (Figure 1C). The bark was then removed using sterile forceps in a large, single piece. The frozen green vascular cambium was then gently scraped from the wood below, into small strips using a sterile scalpel. One hundred milligrams (100 mg) of vascular cambium tissue was finely powdered in microvials containing zirconia beads (BioSpec, Bartlesville, OK, USA) and 550 µL of TRIzol reagent (Invitrogen, Carlsbad, CA, USA) in a MagNA Lyser (Roche, Basel, Switzerland) as described in (Saminathan *et al*., 2014). Finally, RNA was purified using Direct-Zol™ RNA mini-prep kit (Zymo Research, Irvine, CA, USA) and any traces of genomic DNA contamination was removed using enzymatic DNase I treatment. RNA quality and quantity were analyzed using NanoDrop 1100 (NanoDrop, Wilmington, DE, USA) and Agilent 2100 Bioanalyzer (Agilent, Santa Clara, CA, USA).

### cDNA Synthesis and RNA Sequencing

RNA samples for each biological replicate from both treatments (a total of 6 samples) were sequenced at BGI International (http://bgi-international.com/) sequencing platform with standard protocol. Magnetic beads coated with Oligo (dT) were used to isolate mRNA from the total RNA. Using a proprietary fragmentation buffer, the full-length mRNA transcripts were then fragmented into smaller pieces. Next, cDNA was synthesized using random hexamer primers and the mRNA fragments as templates. Short fragments were then purified and resolved with EB buffer for end preparation and single nucleotide A (adenine) addition. The short fragments were then connected with sequencing adapters. For quality control purposes the Agilent 2100 Bioanlyzer and ABI StepOnePlus Real-Time PCR system were used in quantification and qualification of the prepared sample library. Finally, sequencing was performed on the Illumina Hiseq 2000 platform.

### Assembly of Libraries, Data Analysis and Annotation

The raw reads generated from pair-end sequencing are stored in fastq format and usually are contaminated with adapters, unknown or low-quality sequences which were removed by BGI proprietary software “filter_fq”. Once the clean reads were resolved from the raw reads, they were assembled into transcripts using Trinity (http://trinityrnaseq.sourceforge.net/) (Grabherr *et al*., 2011). Three (*Inchworm, Chrysalis*, and *Butterfly*) modules in Trinity were applied sequentially to process raw RNA-seq reads into contigs and full-length transcripts known as unigenes. Inchworm (reference) was further used to assemble the clean reads into unique sequences of transcripts, known as contigs. Chrysalis clustered the Inchworm derived contigs into clusters and constructed complete de Bruijn graphs for each cluster. Butterfly then processed the individual de Bruijn graphs in parallel, tracing the paths that reads and pairs of reads take within the graph, and ultimately reporting full-length transcripts. Butterfly also determined alternatively spliced isoforms of genes. The end result of assembly are full-length transcripts known as unigenes. Open source program BLAST (http://www.blast.ncbi.nlm.nih.gov/Blast.cgi) compares nucleotide, amino acid, or protein sequences to annotated sequence databases and calculates the statistical significance of the homology. The unigenes were aligned with five databases: KEGG (Kyoto Encyclopedia of Genes and Genomes), COG (Clusters of Orthologous Groups), NT (NCBI nucleotide database), NR (NCBI non-redundant protein database) and Swiss-Prot (Protein sequence database). The KEGG database (http://www.genome.jp/kegg/) (Kanehisa *et al*., 2007)was used to perform a systematic analysis of metabolic pathways and function of gene products within a cell. By aligning with the KEGG database the annotated metabolic pathways with which the transcripts (unigenes) correspond were elucidated, allowing insight into the complex biological functions of gene families. The COG database (http://www.ncbi.nlm.nih.gov/COG/) classified orthologous gene products into clusters. COG clusters predicted the possible function of the transcripts. The NT database (http://www.ncbi.nlm.nih.gov/nuccore) is a non-redundant nucleotide database with entries from NCBI’s other databases (GenBank, EMBL, and DDBJ) and offers another way to predict transcript function. Both NCBI’s NR database and the Swiss-Prot (http://www.uniprot.org/uniprot/) annotated protein databases and added additional information about the possible function of the transcripts.

### Gene Ontology and Coding Sequences

Gene ontology (GO) was employed to standardize gene functional classification such as molecular function, cellular component, and biological process. Using the NR database annotation, the Blast2GO program (http://www.blast2go.com/b2ghome) was used to retrieve GO functional classification for all transcripts (Conesa *et al*., 2005). In order to determine the CDS for the transcripts, unigenes were first aligned to the protein databases, listed in order of priority, of NR, Swiss-Prot, KEGG, and COG by using a local blastx (ftp://ftp.ncbi.nlm.nih.gov/blast/executables/release/LATEST/), with significance cutoff value of *e*^<0.00001^, of the unigene sequences. Unigenes with alignments to higher priority databases, for example NR database, were not aligned to lower priority databases. The highest-ranking proteins in the blastx results were used to decide the coding region sequences of unigenes. Results of the blastx alignment used a standard codon table to translate the nucleotide query sequence into a translated amino acid sequence. However, unigenes that could not be aligned to any database were further scanned by ESTScan (Iseli *et al*., 1999), producing nucleotide sequence (5’⟶3’) direction and amino sequence of the predicted coding region.

### Gene Expression Analysis

In order to determine the expression pattern of the unigenes, clean reads were first mapped to unigenes using the program Bowtie2 (v. 2.2.5 http://bowtie-bio.sourceforge.net/bowtie2/index.shtml) (Langmead and Salzberg, 2012). SAM files generated through Bowtie were used with the RSEM (RNA-Seq by Expectation-Maximization) software package (http://deweylab.github.io/RSEM/; v1.2.12) in R (v1.03; http://www.r-project.org/) to measure the expression level of each unigene. RSEM software was used to estimate gene expression levels from RNA-seq data (Li and Dewey, 2011), providing expression data in FPKM (Fragments Per Kilobase of transcript per Million mapped reads) format, which was subsequently used to perform differential gene expression analysis in this study. To detect Differentially Expressed Genes (DEGs) the program NOIseq (http://genome.cshlp.org/content/early/2011/09/07/gr.124321.111) (Tarazona *et al*., 2011) was utilized. In this study winter and spring unigenes with a fold change of ≥ 2 and a probability ≥ 0.8 were considered to be significantly differentially expressed. PCA was accomplished using Princomp function in R.

### Simple Sequence Repeats Analyses

Simple Sequence Repeat (SSR) identification was accomplished with MIcroSAtellite (MISA) software (http://pgrc.ipk-gatersleben.de/misa/misa.html), using the unigenes as input sequences. The identified SSRs which have lengths ≥ 150 bp on both ends of the unigenes were used to design primers. The SSRs which met the selection criteria were used by the software Primer3 (v2.3.4; http://www.onlinedown.net/soft/51549.htm) to design primers. Primers derived from the unigenes were further filtered by removing primers with SSRs within the primer itself and primers aligned to more than one unigene.

### Validation of Wood-forming Genes with RT-qPCR

Based on existing research information, fifteen wood-forming candidate genes correspond to the biosynthesis of cellulose, hemicellulose, or lignin (Dharmawardhana *et al*., 2010; Doering *et al*., 2012; Jia *et al*., 2015; Nieminen *et al*., 2015; Pauly *et al*., 2013; Quang *et al*., 2012), were selected for validation of expression level. Using the local blast utility (ftp://ftp.ncbi.nlm.nih.gov/blast/executables/release/LATEST/) a database of all Paulownia unigenes was created. The mRNA sequences acquired for each of the selected genes were then aligned to the database of all unigenes using the local blastx utility. The unigenes showing maximum homology for each of the genes was selected for two-step RT-qPCR primer design. The software Primer Express v3.01 (Applied Biosystems, Foster City, CA) was used to design primers for the unigenes corresponding to the selected wood forming genes. Complimentary DNA (cDNA) was synthesized from Paulownia vascular cambium total RNA using SuperScript™ II Reverse Transcriptase (Invitrogen, Waltham, Massachusetts, USA) with suggested protocol. FastStart SYBR Green Master Mix (Roche, Grenzacherstrasse, Basel, Switzerland) reagent was used in combination with primers (Table S3) and cDNA. RT-qPCR of three biological replications with no-template control (NTC) involved StepOnePlus Real-Time PCR System (Applied Biosystems, Foster City, CA, USA) and FastStart SYBR Green (Roche). The expression of selected genes was normalized to that of the 18S rRNA gene. Finally, the relative gene expression was calculated by the 2^−ΔΔCt^ method (Livak and Schmittgen, 2001).

## Results and Discussion

Empress tree (*Paulownia tomentosa*), a fast-growing tree species native to China, has been grown for the purpose of timber, pulp, soil protection, and for many other uses. Daylength affects tree growth with a short-day length introduces cessation of growth and growth rate is very high during summer. To ensure survival and productivity, perennial trees in temperate climates utilize cyclical environmental signals, such as daylength and seasonal temperature patterns. The vascular cambium, a lateral meristem found in diverse tree species, is responsible for supporting the radial, woody growth of stems. The vascular cambium consists of meristematic initials that divide over time to produce daughter cells which consequently turn into secondary xylem and secondary phloem of the stem. The daily cumulative temperature is the most important cue for cambial reactivation (Sarvas, 1970), however daylength influences cambial dormancy at the end of the summer and autumn (Heide, 1974). In forest trees, seasonal cues influence several aspects of the secondary growth of tree stems, including cambial activity, wood chemistry, and transition to latewood formation (Jokipii-Lukkari *et al*., 2018). A recent transcriptomic study showed photoperiod as the dominant driver of seasonal gene expression variation in needles of Douglas-fir (Cronn *et al*., 2017). Therefore, it is evident that cambial growth is affected to a greater extent by changes in the ambient temperature and affects overall seasonal growth by tuning thousands of genes reach their annual peak activity during winter dormancy. In this study, we showed how winter and spring seasons modulated the transcriptome and consequently plant growth by studying cambial transcriptome and pathway analysis.

### RNA-Seq and Transcriptome Assembly of Paulownia Cambial Tissue

In order to obtain the candidate genes associated with cambium development of empress tree during seasonal growth, transcriptome sequencing analysis for winter and spring seasons (Figure 1A and 1B) was carried out by collecting cambial tissues from tree twigs (Figure 1C). A total of 305,882,370 (∼29 Gb) raw reads were generated. Removal of adapter sequences, low-quality reads, and ambiguous sequences resulted in 297,049,842 clean reads (Q20 > 97.73%) with an average length of 100 nucleotides. Winter samples generated more raw reads when compared to spring samples (Table 1).

**Table 1.** Summary of the sequence assembly after Illumina sequencing and statistics of contigs and unigenes (n=3). The values are given as Mean±SD from three replications.

|  | Winter Wood | Spring Wood |
| --- | --- | --- |
| Total raw reads | 51,005,253±2,639,904 | 52,359,187±737,515 |
| Total clean reads | 49,766,422±2,349,991 | 50,689,696±1,090,856 |
| Percentage of reads | 97.60±0.52 | 96.64±0.63 |
| Q20 Percentage | 97.96±0.36 | 97.51±0.35 |
| <u>Contigs</u> |  |  |
| Total Number | 129,428±610 | 104,388±1779 |
| Total Length (nt) | 43,730,308±513,387 | 35,501,692±721,304 |
| Mean Length (nt) | 338±3 | 340±1 |
| N50 | 605±7 | 642±4 |
| <u>Unigenes</u> |  |  |
| Total Number | 64,142±1229 | 45,671±1,225 |
| Total Length (nt) | 61,610,800±2,101,797 | 38,465,680±1,457,846 |
| Mean Length (nt) | 960±14 | 842±9 |
| N50 | 1,551±25 | 1,354±22 |

The *de-novo* assembling of clean reads resulted in 129,428 and 104,388 total contigs for winter and spring cambial tissues, respectively. Clustering and assembly of these contigs resulted in 64,142 and 45,671 unigenes for winter and spring tissues with the average length of 960 and 842 nucleotides, respectively. Among the unigenes, all unigenes were sub-classified according to nucleotide length and found that 300 nt category dominated all. The number of unigenes was reduced as the nucleotide length increased from 300 to 3,000 nt (Figure S1). A total of 40,814 genes were greater than 1kb length. Approximately 58,654 unigenes were greater than 500 nucleotides in length. As in most previous studies, the mean length of the contigs (∼340 bp) was shorter than that of the unigenes (>1,000 bp). The paired-end reads resulted in longer unigenes (mean, ∼900 bp) than those reported in previous transcriptome studies on trees (Novaes *et al*., 2008; Wang *et al*., 2010). The mean length of unigenes (900 nucleotides) was less than those in previous studies related to tetraploid and drought (Dong *et al*., 2014b; Xu *et al*., 2014) species of *Paulownia australis* and *P. tomentosa*. Most of our assembled unigenes showed homology to nucleotide sequences to six public nucleotide databases. The unmatched unigenes are most likely to represent Paulownia-specific genes especially related to winter and spring seasons.

### Functional Annotation of Paulownia Cambial Transcriptome

The 61,639 transcripts were annotated by performing a BLAST search of the sequences in six databases namely Nonredundant protein (Nr) database, NCBI non-redundant nucleotide sequence (Nt) database, Swiss-Prot, Kyoto Encyclopedia of Genes and Genomes (KEGG) and Cluster of Orthologous Groups of proteins (COG). Basic Local Alignment Search Tool (BLAST), a sequence similarity search was conducted against the NCBI Nr and Nt database and Swiss-Prot protein database using E-values of less than 10^-5^. The BLAST search of 61,639 unigenes showed similarity of 72.47% to Nr database followed by 69.06% with Nt, 48.97% with Swiss-Prot, 43.17% with KEGG, 29.68% with COG, 53.85% with Interpro and 44.29% with GO (Table S1).

Of the annotated sequences in the non-redundant (Nr) protein database, 39.3% of the mapped unigenes had very significant homology to known sequences (e-value,10–100), 35.1% showed significant homology (10–100, e-value,10–30), and 25.6% showed weak homology (e-value 10–30 to 10–5) (Figure 2A). We also performed the sequence conservation analysis of Paulownia transcripts with proteomes of all sequenced plant species. As depicted in Figure 2B, the E-value distribution analysis of transcripts showed that 47.0% unigenes had a similarity of more than 80%, 49.8% unigenes had a similarity between 40 and 80%, and just 3.2% unigenes had a similarity of less than 40%. We employed a new BLAST operation to study the relationship of Paulownia with other plant species to identify proteins and pathways that would be unique to Paulownia. The sequence conservation analysis of transcripts showed homology to sequences from *Erythranthe guttata* (64.8%), followed by *Vitis vinifera* (6.3%), *Solanum tuberosum* (5.0%), *Solanum lycopersicum* (3.0%), *Theobroma cacao* (2.7%) and others (Figure 2C). *Erythranthe guttata*, a yellow bee-pollinated annual or perennial plant, is a model organism for biological studies. Paulownia transcripts shared strong homology with *Erythranthe* species and this could be due to strong phylogenetic relationship between these two species (Zhao *et al*., 2019). However, transcripts from a drought-related transcriptomic studies of Paulownia (Dong *et al*., 2014b; Xu *et al*., 2014) showed homology to *Vitis vinifera* (45-48%). This shift in homology could be due to the selection of abiotic seasonal tissue for cambial transcriptome study.

**Figure 2.**
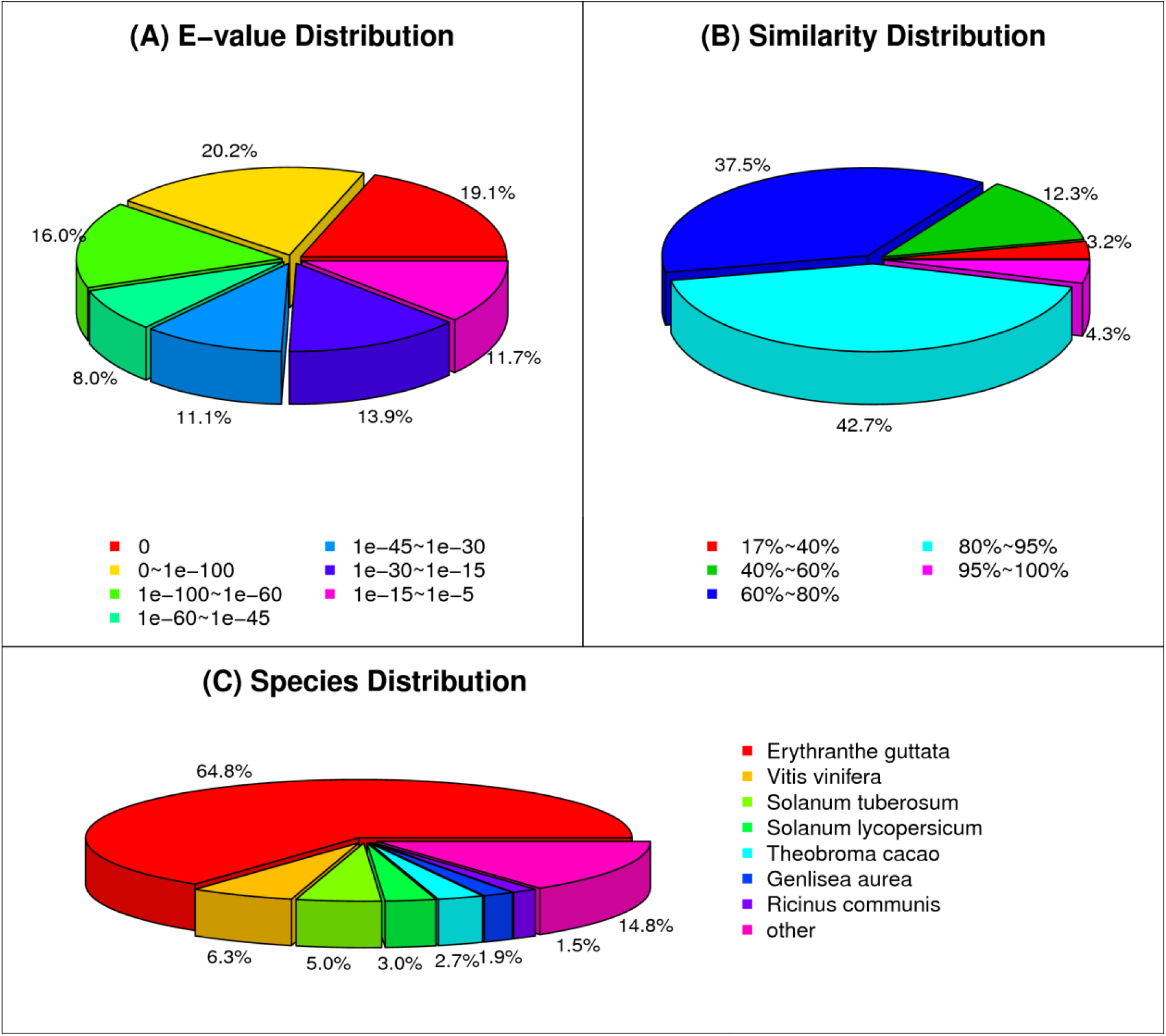
Statistics of homology search of unigenes against Non-redundant (NR) protein database. Distribution of top BLASTX hits with cut-off *e*-value of <1.0 x 10^-5^ (A), similarity (B), and species distributions (C) of all unigenes.

The analysis of 43,780 transcripts in COG database classified them into 25 protein families participated in transcription, replication and recombination, posttranslational modification, signal transduction, and so on. However, the cluster for general function prediction (7,998) represented the largest group, followed by transcription (4,017), replication, recombination and repair (3,451), posttranslational modification, protein turnover and chaperones (3,414), signal transduction mechanisms (3,158), translation, ribosomal structure and biogenesis (3,055), carbohydrate transport and metabolism (2,776), amino acid transport and metabolism (1,874), cell wall/membrane/envelope biogenesis (1,629), energy production and conversion (1,563), cell cycle control, cell division, and chromosome partitioning (1,356). In contrast, only a few unigenes were assigned to extracellular structure and nuclear structure (17 and 4 unigenes, respectively). Importantly, many unigenes have been assigned to a wide range of COG classifications (Figure 3), indicating that a wide diversity of transcripts involved in wood formation as in Chinese fir (Qiu *et al*., 2018).

**Figure 3.**
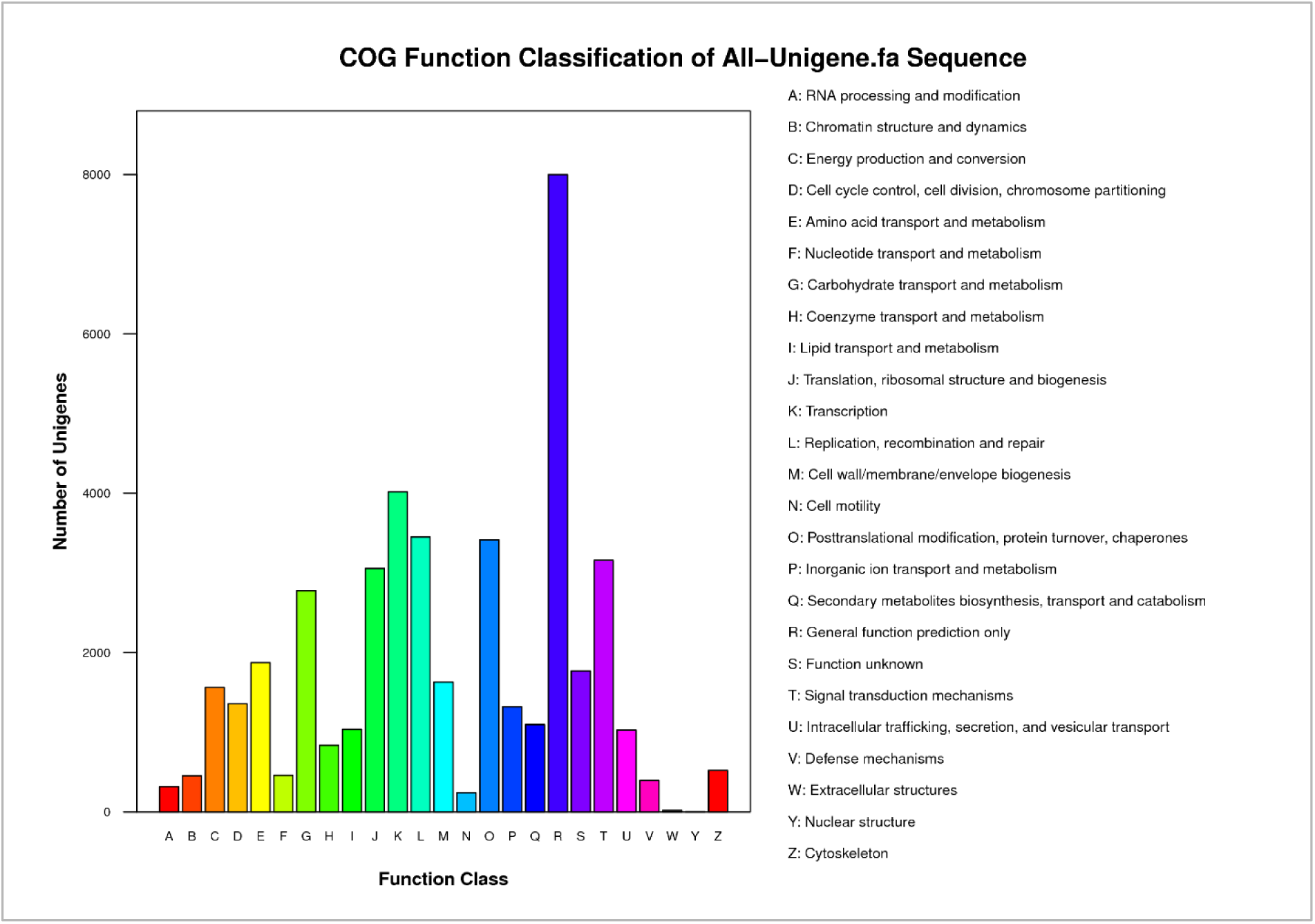
Histogram representation of clusters of orthologous groups (COG). The horizontal coordinates are function classes of COG, and the vertical coordinates are numbers of unigenes in one class. The notation on the right is the full name of the functions in X-axis. Histogram representation of classification of the clusters of orthologous groups (COG) for the total aligned 43,780 unigenes (53.43%) into 25 functional groups.

The Gene Ontology (GO) classification classified 42,588 out of 62,639 transcripts into ontologies related to molecular functions, cellular components and biological processes, allowing a coherent annotation of transcripts (Figure 4). We identified significantly higher number of transcripts (19,418) involved in metabolic process and 18,047 related to cellular processes (18,047) when compared to others such as rhythmic processes (number). The most represented category for cellular components was cells (GO: 0005623; 18186 genes) followed by organelle (GO:0043226; 14053 genes). But for molecular functions, the catalytic activity (GO: 0003824; 17403 genes) was the most represented GO term followed by the binding activity (GO: 0005488; 15485 genes). Genes and pathways putatively responsible for dormant winter and active spring growth in Paulownia were identified in this study. In *Populus, PtrHB7*, a class III HD-Zip gene, is known to play a critical role in regulation of vascular cambium differentiation (Zhu *et al*., 2013) and homeobox gene *ARBORKNOX1* regulates the shoot apical meristem and the vascular cambium (Groover *et al*., 2006). In our study, Unigene2201, Unigene3374, and Unigene4121 which were downregulated belong to GO:0005488 (molecular function: binding) and are homologs of *KNOX* gene. *KNOX* family gene *KNAT7* negatively regulates secondary wall formation in *Arabidopsis* and *Populus* (Li *et al*., 2012). Since *KNAT7* is a negative regulator of secondary wall biosynthesis, these Paulownia homologs might positively regulate cambium growth during active spring season.

**Figure 4.**
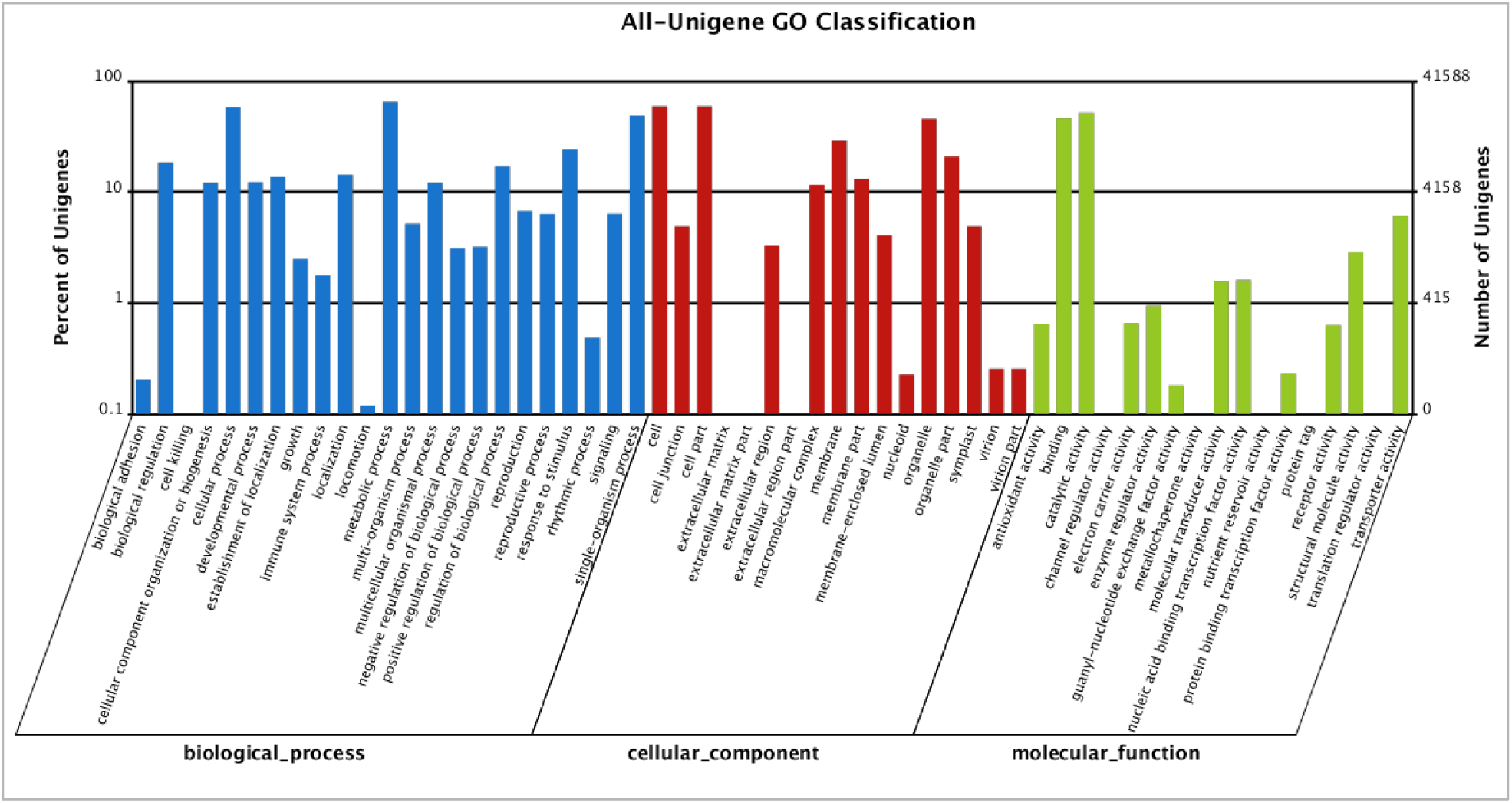
GO classification analysis of unigenes. GO functions is showed in X-axis. The right Y-axis shows the number of genes which have the GO function, and the left Y-axis shows the percentage. Unigenes in winter and spring season are classified into biological processes, cellular component and molecular function. In total 41,588 (50.76% of all unigenes) were assigned to 48 GO categories.

In order to categorize the different biochemical pathways that the annotated unigenes were associated with, we assigned the EC numbers in the KEGG pathways. KEGG annotation yielded a total of 35,471 (57.5%) unigenes that were mapped to 128 KEGG pathways. The top eight KEGG enriched pathways were metabolic pathways 7943 (22.39%; ko01100) biosynthesis of secondary metabolites 3768 (10.62%; ko01110), and plant-pathogen interaction 1852 (5.22%; ko04626), plant hormone signal transduction 1717 (4.84%; ko04075), spliceosome 1462 (4.12%; ko03040), ribosome 1179 (3.32%; ko03010), RNA transport 1165 (3.28%; ko03013) and protein processing in endoplasmic reticulum 1120 (3.16%; ko04141) (Table S2). With the help of KEGG database, we could further analyze the metabolic pathways and functions of gene products, which help in studying the complex biological behaviors of genes. Most representative unigenes were annotated to metabolic pathways, biosynthesis of secondary metabolites, plant-pathogen interaction, plant hormone signaling, spliceosome, and phenylpropanoid biosynthesis using the KEGG database, lead us conclude that most of the genes identified in this study are involved in cambial differentiation and wood formation.

### Transcriptional Profiling of Cambial Tissues in Winter and Spring

A total of 10,099 (12.33%) transcripts were found to be significantly differentially expressed between two tissue samples. Of these differentially expressed genes (DEGs), 2,688 (26.61%) transcripts were found to be upregulated (>1.6 fold) in spring season, whereas 7,411 (73.39%) were downregulated (<–1.6 fold) when compared to winter season (Figure S2). Hierarchical clustering of the DEGs identified in winter and spring conditions led to the detailed overall structure of clustering. This indirectly indicated that more genes were upregulated, active and required during the senescent winter season to keep tissues dormant. Out of 2,688 genes, top 20 genes with log2Fold change >8.00 are summarized in Table 2. This included APC/C cyclosome complex, phosphoenolpyryvate carboxy kinase, different classes of heat shock proteins, actin depolymerization factor, anaphase-promoting complex subunit (>12-fold expression), etc. Similarly, many key genes including synthases such as galactinol synthase (<– 12-fold expression), rosmarinate synthase, and valencene synthase, kinases such as receptor-like protein kinase, serine/threonine protein kinase, and CBL-interacting protein kinase, and hormone regulated genes such as auxin efflux carrier family protein and ethylene-responsive transcription factor were downregulated (Table 3). The cell cycle, being one of the most important biological processes in the cambial zone, plays central role in regulating the growth and development of organisms including plants. The anaphase-promoting complex/cyclosome (APC/C; homolog in our study Unigene8688), a well-known ubiquitin ligase, acts to accomplish basic cell-cycle control. The APC/C must be turned off at the end of G1 phase to allow the S phase cyclins to accumulate and cells to begin DNA replication (Pines, 2011). This is very key during spring season for cell multiplication and growth. The *Cyclin U2* (Unigene22553), one of the major cyclins involved in cell cycle control, like cyclins A and B on maximum gene expression in poplar cambium zone (Hertzberg *et al*., 2001) was upregulated in Paulownia. The high abundance of cyclin transcripts in active cambium during spring season also reflected a positive correlation between cambium cell division and key cell cycle gene expression.

**Table 2.**
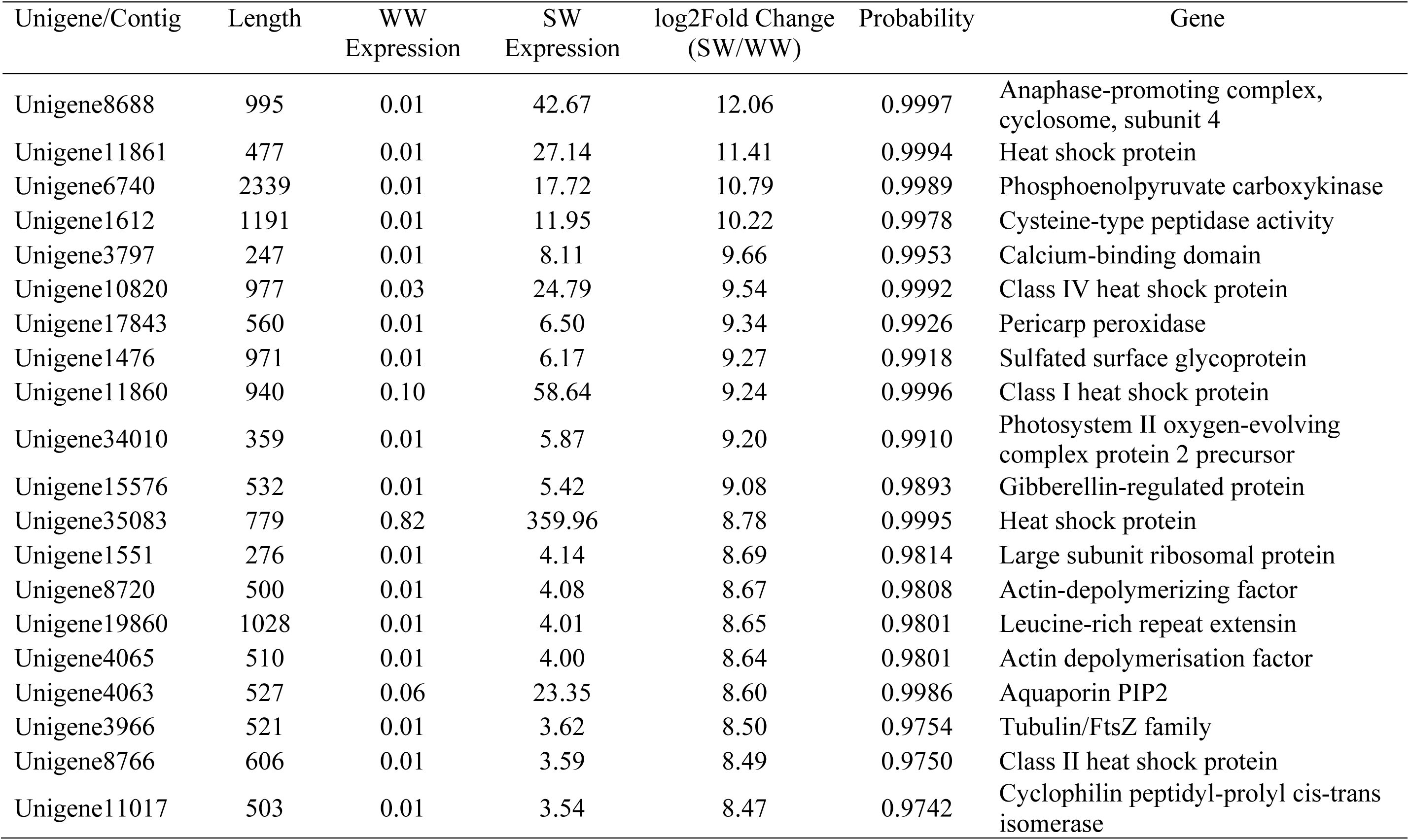
List of top 20 upregulated known genes

**Table 3.**
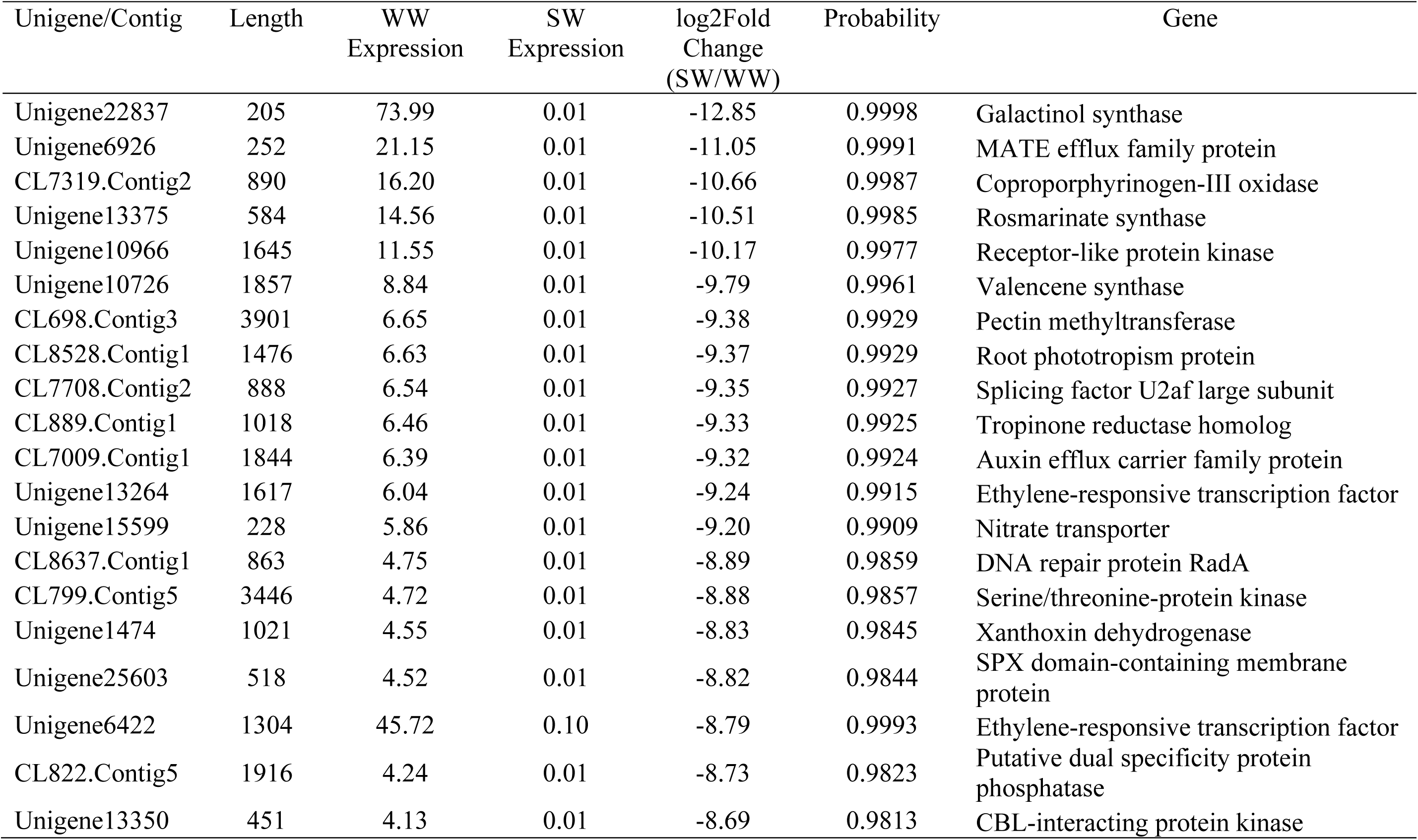
List of top 20 downregulated genes

As shown in Figure 5, GO analysis indicated that most of the DEGs for biological process were involved in the metabolic process (1,016), cellular process (890), and response to stimulus (350). GO cellular component analysis revealed the cell (824), cell part (789), membrane (436) and organelle (399) enriched the most DEGs. Meanwhile, GO molecular function analysis showed that the DEGs predominantly contributed to catalytic activity (836) followed by transporter activity and structural molecule activity. KEGG enrichment analysis of DEG showed that these genes were involved in various pathways in Paulownia plant during seasonal changes (Table 4). Most of the DEGs were enriched in metabolic pathways (ko01100; 1387), biosynthesis of secondary metabolites (ko01110; 827), and plant hormone signal transduction (ko04075; 320). It was also found that starch and sucrose metabolism (ko00500; 172; Figure S3) and phenylpropanoid biosynthesis (ko00940; 106; Figure S4), which correspond to the production of several key wood forming genes, were within the top 25 most DEG enriched KEGG pathways (Table 4). Nineteen (Unigene11539, Unigene12164, Unigene12788, Unigene16018, Unigene17615, Unigene18048, Unigene18594, Unigene18926, Unigene22808, Unigene24462, Unigene24837, Unigene3634, Unigene4753, Unigene6221, Unigene891, and Unigene9670 (K00430), Unigene16856 (K11188), Unigene22599 and Unigene25305 (K03782)) out of 497 unigenes (total unigenes) involved in lignin synthesis in the phenylpropanoid biosynthesis pathway (Ko00940) were identified and differentially regulated (Figure S4). Lignin plays a vital role in keeping the structural integrity of the cell wall, and protecting plants from pathogens (Hu *et al*., 2008) as well as a main component of wood. Of these 19, different types of peroxidases (Unigene11539, Unigene18926, Unigene4753 and Unigene6221) were upregulated during winter. Recently, a notable remodeling of the transcriptome was reported in Norway Spruce where monolignol biosynthesis genes showed high expression during the period of secondary cell wall formation as well as a second peak in midwinter. Interestingly, this midwinter peak expression did not trigger lignin deposition. (Jokipii-Lukkari *et al*., 2018). These genes could be preparing for the biosynthesis and distribution of guaiacyl (G), p-hydroxyl phenol (H) and syringyl (S) lignin in developing biomass as soon as the onset of Spring.

**Table 4.** Top 25 DEG enriched KEGG pathways.

| Pathway | Number of<br>DEGs genes | *p*-value | Pathway ID |
| --- | --- | --- | --- |
| Metabolic pathways | 1387 | 1.32E-12 | ko01100 |
| Biosynthesis of secondary metabolites | 827 | 7.55E-10 | ko01110 |
| Plant hormone signal transduction | 320 | 1.58E-05 | ko04075 |
| Plant-pathogen interaction | 266 | 0.2430222 | ko04626 |
| Ribosome | 204 | 0.200625 | ko03010 |
| Spliceosome | 178 | 0.9468082 | ko03040 |
| Starch and sucrose metabolism | 172 | 1.22E-06 | ko00500 |
| Protein processing in E.R. | 162 | 0.3255727 | ko04141 |
| Carbon metabolism | 161 | 0.04321163 | ko01200 |
| RNA transport | 151 | 0.986695 | ko03013 |
| Glycerophospholipid metabolism | 138 | 0.1244536 | ko00564 |
| Endocytosis | 134 | 0.6458048 | ko04144 |
| Biosynthesis of amino acids | 123 | 0.7258124 | ko01230 |
| Glycolysis / Gluconeogenesis | 115 | 5.48E-05 | ko00010 |
| Phenylpropanoid biosynthesis | 106 | 1.50E-06 | ko00940 |
| Circadian rhythm - plant | 105 | 5.10E-09 | ko04712 |
| Ether lipid metabolism | 104 | 0.02664109 | ko00565 |
| Ubiquitin mediated proteolysis | 99 | 0.7266876 | ko04120 |
| Pentose and glucuronate interconversions | 93 | 1.57E-05 | ko00040 |
| Purine metabolism | 92 | 0.9911015 | ko00230 |
| Amino sugar and nucleotide sugar metabolism | 86 | 0.006516035 | ko00520 |
| Pyrimidine metabolism | 78 | 0.992583 | ko00240 |
| mRNA surveillance pathway | 77 | 0.9938776 | ko03015 |
| Flavonoid biosynthesis | 70 | 8.18E-07 | ko00941 |
| RNA degradation | 68 | 0.940714 | ko03018 |

**Figure 5.**
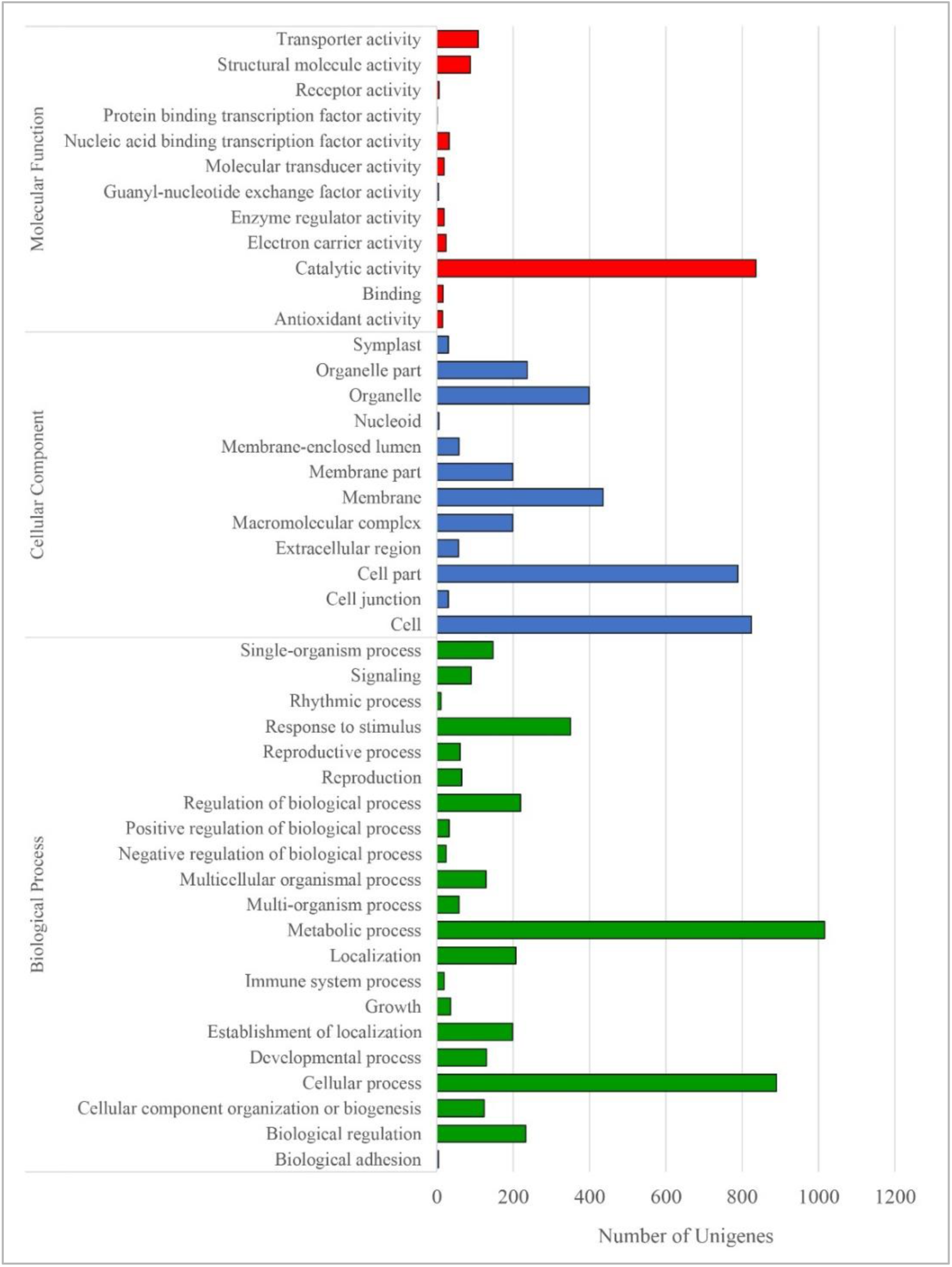
GO function analysis of the differentially expressed genes. GO function analysis results for the differentially expressed genes in cambial tissues due to winter and spring seasons into biological processes, cellular component and molecular function.

Out of 61,639 annotated unigenes, 58,324 unigenes were matched to known genes using blastx and 1,708 unigenes were matched to coding sequences using ESTscan. Most unigenes population were ranked from 200-3000 nucleotides in length (Figure S7). Most of the unigenes (34,000) were about 200 nt in length for coding sequences. There were no translated peptides beyond 2,500 sequences. In case of ESTScan method, most of the unigenes from 200-3000 nt in length were translated into protein sequences in the range of 200 to 1,100. This difference could be due to extra sequences in the full-length cDNAs than protein coding sequences.

### Expression of Lignocellulosic Pathway Genes and Their Validation

Wood, the secondary xylem, is produced from the activity of vascular cambium that is composed of two meristematic initials: fusiform initials and ray initials (Mauseth, 1988) with the sequential developmental process including differentiation of vascular cambium cells into secondary xylem mother cells, cell expansion, and massive deposition of secondary walls where a number of genes involved in vascular tissue differentiation and secondary wall biosynthesis (Zhong and Ye, 2015). When the wood compression starts, the expression of a number of genes involved in synthesis of lignocellulosic components (cellulose, hemicellulose and lignin) and lignans was upregulated in maritime pine (Villalobos *et al*., 2012). In addition onset of wood formation undergoes three periods: winter shrinkage, spring rehydration (32-47 days) and summer transpiration in the stem (Turcotte *et al*., 2009).

In order to explore the roles of cell wall- and hormone related genes for the seasonal cues, fifteen candidate genes were identified from previous studies (Table 5). They are involved in cellulose (*CesA1, CesA3, CesA6* (Djerbi *et al*., 2004)), hemicellulose (*CSLC4* (Davis *et al*., 2010), *FUT1* (Perrin *et al*., 1999; Vanzin *et al*., 2002), *AXY4* (Gille *et al*., 2011), *GATL1* (Kong *et al*., 2009), *IRX10 (Hörnblad et al*., *2013), ESKIMO1* (Lefebvre *et al*., 2011)), lignin (*4CL (Hu et al*., *1999), CCR1* (Goujon *et al*., 2003), *CAD1* (Bouvier d’Yvoire *et al*., 2013)), auxin (*TAA1* (Stepanova *et al*., 2008), *YUC1* (Cheng *et al*., 2006; Won *et al*., 2011)) and cytokinin (*IPT1* (Immanen *et al*., 2016)) synthesis/pathways. RT-qPCR was employed to study the expression of these wood formation genes in Paulownia during winter and spring seasons (Figure 6).

**Table 5:**
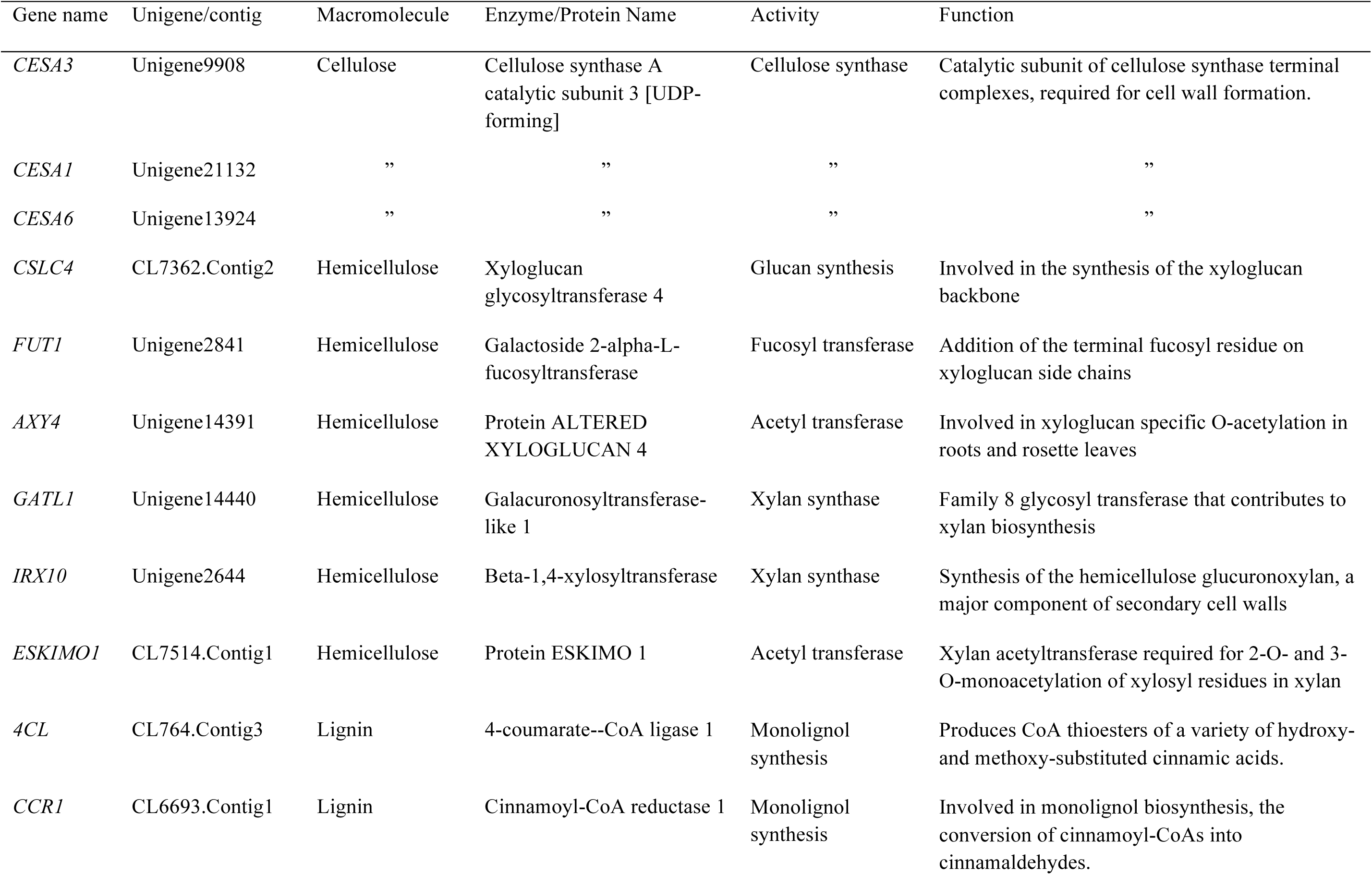
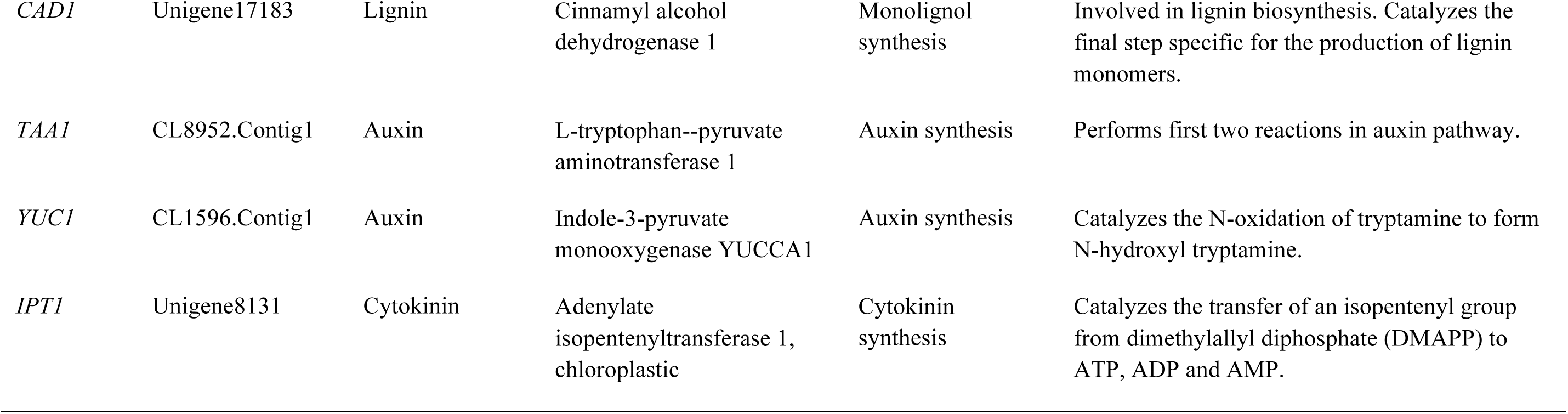
Wood-Forming Genes Selected for RT-qPCR Expression Validation

**Figure 6.**
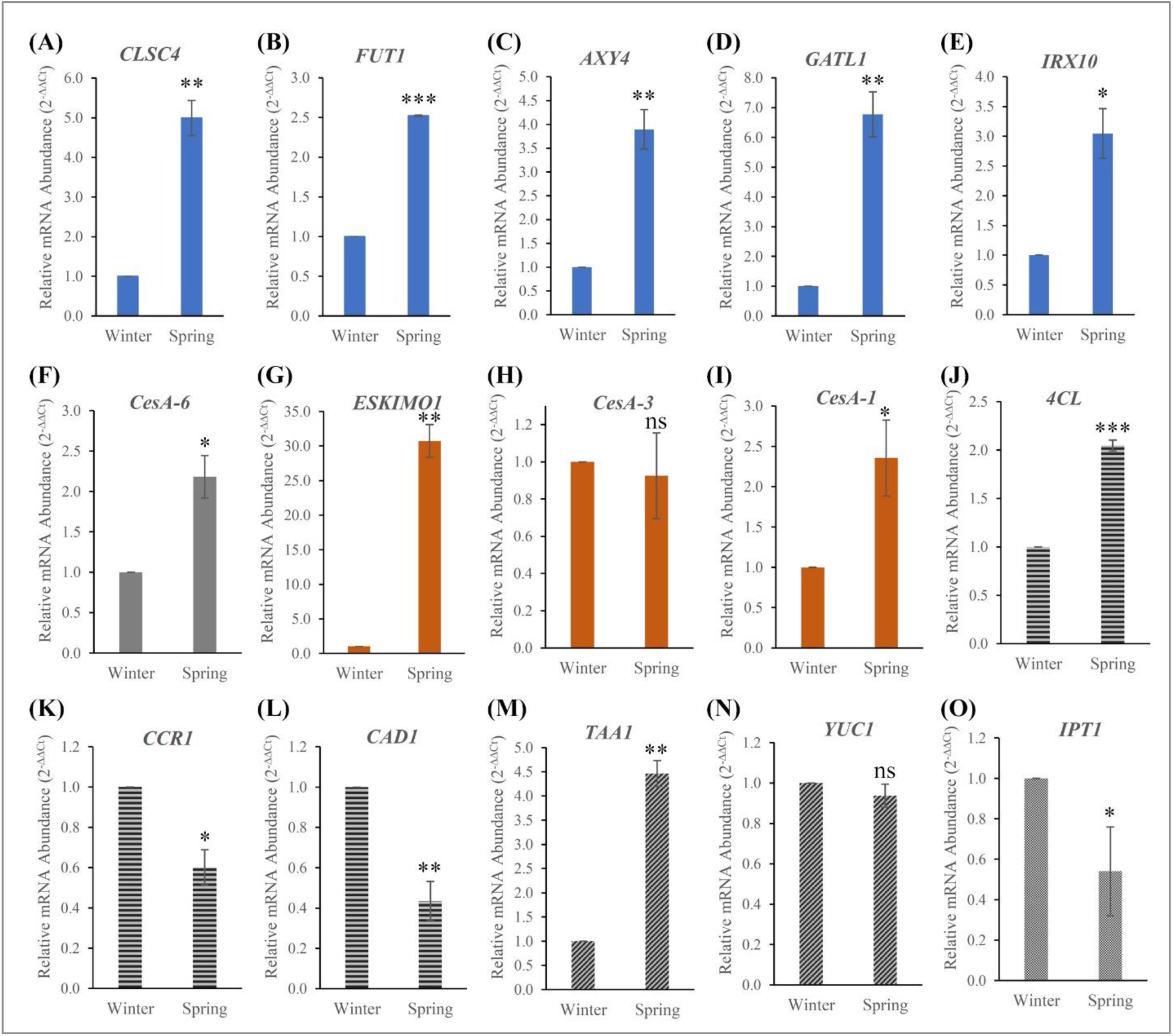
Relative mRNA expression of key genes involved in winter and spring seasons. Expression of genes involved in cell wall synthesis (*CESA3, CESA1, CESA6, CSLC4, FUT1, AXY4, GATL1, IRX10, ESKIMO1, 4CL, CCR1, CAD1*) and hormone synthesis (*TAA1, YUC1, IPT1*). Expression quantity of the calibrator sample (winter tissue) was set to 1. Data are the mean ± SD. Student’s *t*-test was used to compare significant changes in spring tissues compared to winter tissues. *, *p* <0.1; **, *p* < 0.01; ***, *p* < 0.001; ns; no significance.

Cellulose is synthesized in plant cell walls by large membrane-bound protein complexes proposed to contain several copies of the catalytic subunit of the cellulose synthase, *CesA*. Here, we found *CesA1* and *CesA6* were upregulated during spring while *CesA3* was moderately downregulated during winter season. In hybrid aspen, expression analyses of CesA family showed specific location in normal wood undergoing xylogenesis, while *PttCesA2*, seems to be activated on the opposite side of a tension wood (Djerbi *et al*., 2004). However, in Arabidopsis, the expression levels of the three primary cell wall genes (*AtCesA2, AtCesA5, AtCesA6*) was increased, but not *AtCesA3*, AtCesA9 or secondary cell wall *AtCesA7* (Hu *et al*., 2018). Our results along with these studies indicated that the expression of major primary wall CesA genes to accelerate primary wall CesA complex.

Several proteins encoded by the cellulose synthase□like (*CSL*) gene family including CSLA proteins, which synthesize β□(1→4)□linked mannans, and CSLC proteins, which are thought to synthesize the β□(1→4)□linked glucan backbone of xyloglucan are known to be involved in the synthesis of cell□wall polysaccharides (Davis *et al*., 2010). Higher expression of CLSC4 in Paulownia during spring season indicated that it might involve cellulose synthesis. The fucosyltransferase (FUT1) is an enzyme that transfers an L-fucose sugar from a GDP-fucose (guanosine diphosphate-fucose) donor substrate to an acceptor substrate. The Arabidopsis *mur1* (AtFUT1) mutant study (Vanzin *et al*., 2002) exhibited a dwarf growth habit and decreased wall strength indicating indispensable role of FUT1 function in wood formation. Another key gene family of O-acetyl substituents seems to be very important for various plant tissues and during plant development (Liners *et al*., 1994), suggesting an important functional role in the plant. Mutants lacking *AXY4* transcript resulted in a complete lack of O-acetyl substituents on xyloglucan in several tissues, except seeds (Gille *et al*., 2011). Biosynthesis of xylan in woody plants is a major pathway for plant biomass. Populus genes *PdGATL1*.*1* and *PdGATL1*.*2*, the closest orthologs to the Arabidopsis *PARVUS/GATL1* gene, have been shown to be important for xylan synthesis, but may also have role(s) in the synthesis of other wall polymers (Kong *et al*., 2009). The expression of *GATL1* homolog in Paulownia was six-fold (Figure 6) in spring season implying more xylan biosynthesis. Collapsed xylem phenotypes of Arabidopsis (Turner and Somerville, 1997) and *Physcomitrella patens* (Hörnblad *et al*., 2013) mutants (*irx10*) identified mutants deficient in cellulose deposition in the secondary cell wall due to lack of synthesis of the glucuronoxylan. Acetyl transferases are involved cellulose biosynthesis in plants. In Arabidopsis, the *ESKIMO1* (*ESK1*) gene has been described for multiple roles and mutants of which (*esk1*) indicated that *ESK1* is necessary for the synthesis of functional xylem vessels towards laying down of secondary cell wall components (Lefebvre *et al*., 2011). Our gene expression along with RNAseq of cellulose and hemicellulose indicated that all these genes were highly expressed during spring season to prop complete wood formation.

Trees have extreme needs for both structural support and water transport and 15 to 36% of the dry weight of wood is lignin (Sarkanan and Hergeht, 1971). Since lignin limits the use of wood for fiber, chemical, and energy production, lignin is therefore one of the world’s most abundant natural polymers, along with cellulose and chitin. It has been shown that *PAL* or *4CL* (*4□coumarate:coenzyme A ligase*) was strongly downregulated indicating lower lignin content with a preferred reduced content in G units and increase cellulose in aspen (Hu *et al*., 1999). However, we found upregulation of *4CL* in Paulownia (Figure 6). It is reported that Paulownia spp. and hybrids showed more Klason lignin 22.9–27.8% whereas aspen exhibited 19.3% (García *et al*., 2011) relating positive correlation with increased *4CL* transcripts in Paulownia from RT-qPCR results. Cinnamoyl CoA reductase (CCR) is the first enzyme in monolignols synthesis. Arabidopsis (*atccr1*) mutants were severely downregulated and had 50% decrease in lignin content accompanied by changes in lignin composition and structure implying *CCR1* was a positive regulator (Goujon *et al*., 2003). However, it showed downregulation in Paulownia during spring season. Disrupting the *Brachypodium Cinnamyl Alcohol Dehydrogenase 1* gene (*BdCAD1*) leads to altered lignification and improved saccharification (Bouvier d’Yvoire *et al*., 2013). Another study in poplar tree suggested that downregulating *CAD1* is a promising strategy for improving lignocellulosic biomass (Van Acker *et al*., 2017). Lignin is the major phenolic polymer in plant secondary cell walls and is polymerized from monomeric subunits, the monolignols (Yan *et al*., 2019). Thr BiFC biochemical assay showed molecular interaction of *PtrCAD1*/*PtrCCR2* homo- and heterodimer formation. The downregulation of CCR1 and CAD1 in Paulownia indicates the improvement of lignocellulosic biomass. Altogether, the lignocellulosic pathway genes regulate the components of cellulose, hemicellulose and lignin in appropriate ratios as indicated earlier repression of lignin biosynthesis promotes cellulose accumulation and growth in aspen tree (Hu *et al*., 1999).

### Analysis of Hormone-Specific Genes and Their Validation

Auxins, morphogen-like plant-growth regulators, with some play a key role in regulating wood formation through its effects on cambial activity and xylem development (Sundberg *et al*., 2000). It is required for maintaining the cambium in a meristematic state as depleting the cambium of auxin leads to differentiation of cambial cells to axial parenchyma cells (Savidge, 1983). Cytokinins, on the other hand, have a well-established function in cell division during growth and development and they are called central regulators of cambial activity (Matsumoto-Kitano *et al*., 2008). The interaction between auxin and cytokinin seems to be essential for induction of phenylalanine ammonialyase activity in support of lignification (Bevan and Northcote, 1979). *TAA1*, which performs first two reactions in auxin pathway, is a Trp aminotransferase that converts Trp to IPA in the IPA auxin biosynthesis branch in Arabidopsis (Won *et al*., 2011). Higher order mutants in *TAA1* showed auxin-related multiple phenotypes. Later, it was identified that *TAA1* gene was essential for hormone crosstalk with ethylene for plant development (Stepanova *et al*., 2008). Later, new putative function of IAA production via IPyA and transport was identified which was newly postulated (Stepanova *et al*., 2011).

Another group of auxin biosynthesis gene family, *YUCCA* flavin monooxygenases, controls the formation of floral organs and vascular tissues in Arabidopsis (Cheng *et al*., 2006). When *TAA* family of amino transferases converts tryptophan to indole-3-pyruvate (IPA) and that the *YUCCA* (*YUC*) family participates in converting IPA to IAA, the main auxin in plants (Won *et al*., 2011). In addition, the authors found that *YUC* and *TAA* work genetically in the same pathway and that *YUC* is downstream of *TAA*. From our transcriptome and gene expression studies, we observed *TAA1* was strongly expressed with no obvious difference in *YUC1* expression during spring season. Different unigenes involved in auxin biosynthesis are given in tryptophan pathway for cell enlargement and plant growth (Figure S5).

In Arabidopsis, cambial activity responded to small changes in cytokinin levels indicating that cytokinins are central regulators of cambium activity (Matsumoto-Kitano *et al*., 2008). Isopentenyltransferase, the rate limiting step of cytokinin biosynthesis, is an important enzyme playing key roles in meristem maintenance and organ development. Arabidopsis quadruple mutants lacking *AtIPT1, AtIPT3, AtIPT5*, and *AtIPT7* were unable to form cambium and showed reduced thickening of the root and stem, thought single mutant *atipt3* showing moderately decreased levels of cytokinins without any other recognizable morphological changes. Similarly, increased cytokinin biosynthesis stimulates the cambial cell division rate and increases the production of trunk biomass in transgenic Populus trees (Immanen *et al*., 2016). Surprisingly, *IPT1* expression was high in winter and moderately reduced in spring indicating two possibilities. Cytokinin pathway with *IPT1* role might have been active during mid-winter (end of March). Alternatively, there could be many other members in *IPT* family that complement the function of cambial development. Auxin and cytokinin display distinct distribution profiles across the cambium and elevated cytokinin content leads to an increased cambial auxin concentration (Immanen *et al*., 2016). Together, it is very interesting to see the interaction of lignocellulosic pathways genes along with major hormone-regulated genes and their crosstalks to maintain the balance of cambial activities for quality wood formation with alternative seasonal changes (Figure S6).

### Analysis of Simple Sequence Repeats (SSRs)

SSR markers are very useful for multiple applications in plant genetics because of their co-dominance, high level of polymorphism, multi-allelic variance, and abundance, and cross-species transferability (Barbara *et al*., 2007; Powell *et al*., 1996). In the present study, SSR were identified utilizing the transcriptome of paulownia cambial tissues because EST-SSR markers have a relatively higher transferability than genomic SSRs (Varshney *et al*., 2005). Recent studies showed that abundant EST-SSRs from RNA-seq have agronomic potential and constitute a scientific basis for future studies on the identification, classification, molecular verification and innovation of germplasms in hawthorn and Lei bamboo (Cai *et al*., 2019; Ma *et al*., 2019).

We identified 11,338 SSRs from the annotated 61,639 unigenes. We detected 3,036 mononucleotides, 5492 dinucleotides, 2493 trinucleotides, 204 tetranucleotides, 194 pentanucleotides, and 344 hexanucleotides motifs (Figure 7; Table S4). Among the dinucleotide and trinucleotide SSRs, AG/CT repeats represented 2,997 SSRs, and AAG/CTT repeats represented 582 SSRs. In mononucleotide, dinucleotide, trinucleotide, quadnucleotide, pentanucleotide, and hexanucleotide repeat categories, the occurrences of repeats were twelve, six, five, five, four and four, respectively (Table S4). Finally, 6,773 oligonucleotide pairs were generated for these identified SSR markers (Table S5). SSRs and SNPs are the most useful and robust molecular markers for genetics and plant breeding applications (Hiremath *et al*., 2012). This study provided a set of SSR markers that could be used, for example, in diversity analysis of Paulownia species. In addition, Paulownia tree breeding programs will benefit from the availability of these SSR markers identified from our RNAseq data. Mononucleotide SSRs would be excluded because of the frequent homopolymer errors found in sequencing data and less polymorphism, dinucleotides (46.6%) and trinucleotides (21.2%) contributed most in Paulownia. This is consistent with the EST-SSRs distributions reported in other plant species (Ahn *et al*., 2013; Wang *et al*., 2014). In plants, SNPs are predominantly beneficial in the construction of high-resolution genetic maps, positional cloning, marker assisted selection (MAS) of important genes, genome wide linkage disequilibrium associate analysis, and species origin, relationship and evolutionary studies (Shahinnia and Sayed-Tabatabaei, 2009).

## Conclusion

Paulownia is a fast growing, multipurpose timber tree suitable for use as a dedicated lignocellulosic bioenergy crop. In order to understand the genes involved in formation of woody biomass related to seasonal cues, a *de novo* transcriptome study was conducted on vascular cambium tissue from senescent winter vascular cambium tissue and actively growing spring vascular cambium tissue. To the best of our knowledge, this is the first transcriptome-based study on *P. elongata*, as well as the first transcriptome study performed on Paulownia vascular cambium tissue focusing seasonal difference. A set of transcripts was specifically expressed in two different tissues. The transcript abundance data confirms the differential pattern of expression of cellulosic, hemicellulosic, lignin biosynthesis specific, and hormone pathway specific genes. By analyzing the transcriptome from two different temporal treatments (winter and spring), representing two distinct physiological states of the plant, DEGs were identified from both treatments. Cell division is one of the key process taking place in the cambial zone and majority of the cell cycle genes were upregulated during the active stage. Onset of cambial activity began between the end of March and the beginning of April as the increased vacuolization of meristematic cells and the mitotic activity occur. However, our study showed more genes were downregulated in spring season remain to be answered. Overall, results of this study will be useful for future research regarding wood formation in Paulownia and other trees.

## Figure Legends

Figure 7. Simple Sequence Repeat (SSR) marker variation statistics. Number of motifs are given against each repeated nucleotide categories from mono-nucleotides to hexa-nucleotides.

